# Limitations of inferring antiviral efficacy of interfering particles from observational natural histories

**DOI:** 10.64898/2026.03.11.708863

**Authors:** Neha Khetan, Gustavo Vasen, Davey Smith, Leor Weinberger

**Affiliations:** Dept. of Biochemistry and Biophysics, University of California, San Francisco USA; University of San Andres, Buenos Aires, Argentina; Dept. of Medicine, Center for AIDS Research, University of California, San Diego USA; Autonomous Therapeutics Inc. Rockville MD, USA; Dept. of Cell & Systems Biology, University of Miami, Miller School of Medicine USA

## Abstract

Recently Hariharan et al.^2^ reported naturally arising defective HIV genomes capable of conditional replication and interference in humans. While this work makes an important contribution to the field of therapeutic interfering particles (TIPs), particularly with respect to safety and tolerability, it also raises fundamental issues regarding: (i) whether the presented data constitute a valid test of, or support conclusions about, the therapeutic potential of TIPs and (ii) technical issues pertaining to the reported basic reproductive numbers (R_0_). Hariharan et al. conclude that the findings “raise concerns about the effectiveness of TIPs.” However, the data presented do not constitute a valid test of therapeutic efficacy. Here, we (i) clarify that post-hoc observational natural history cannot adjudicate the success or failure of an intervention, (ii) show new analysis highlighting how the reported R_0_ measurements are internally inconsistent with the within-host viral dynamics reported, and (iii) explain that alternative, well-established mechanisms remain sufficient to explain the reported observations.

## Introduction

Hariharan et al. demonstrate that defective HIV genomes can persist, diversify, and conditionally propagate in humans over extended periods without causing detectable toxicity, accelerated disease, or novel pathogenic effects. While the study was not designed to assess safety, this observation supports the biological tolerability of defective HIV genomes *in vivo*—a central regulatory concern for TIP-based therapeutics. The absence of adverse clinical consequences despite long-term persistence of these particles in people with HIV argues that sub-genomic defective viral particles appear tolerated in persons with HIV.

At a fundamental level, however, the authors’ stated conclusions about efficacy rely on observational interpretation rather than interventional evidence^*3*^. A well-established body of literature^4,5^ recognizes that adjudicating therapeutic efficacy from observational natural history is invalid; accordingly, many journals restrict any such interpretation to prospective randomized interventional trials.

Another major source of ambiguity is that all observations in the Hariharan et al. study were made in individuals receiving antiretroviral therapy (ART). However, defective HIV genomes encode the same HIV target enzymes (e.g., reverse transcriptase) as wild-type virus and are necessarily susceptible to antiretroviral drugs. Moreover, given that defective genomes depend on active HIV replication, ART should suppress defective genomes at least as effectively as replication-competent full-length virus—plausibly more so. In this setting, the study design does not permit inference about the therapeutic efficacy of defective interfering particles (DIPs) or TIPs.

Consequently, attributing the absence of strong therapeutic benefit specifically to defective genomes may not be justified when ART itself failed to fully suppress viremia in these persons. The same reasoning could lead some to raise concerns about the therapeutic effectiveness of ART, which would be unwarranted.

## Results

A major conundrum is raised by the reported R_0_ values. Hariharan et al. report an effective R_0_ for wild-type HIV below unity (R_0_ < 1) under their particular assay conditions. By definition, an R_0_ < 1 implies inevitable extinction of the virus, which is mathematically incompatible with the persistent, sustained HIV viremia (i.e., viral set point) observed in the studied individuals (P1 and P2). No within-host equilibrium can exist if the reproductive number of the virus is below unity and as demonstrated from canonical within host models (**Fig. 1**), there is no combination of R_0_ values for a DIP (or TIP) and HIV that can produce sustained viral set points when HIV R_0_ < 1. Consequently, the assay results appear internally inconsistent with the within-host viral dynamics and the reported R_0_ estimates may not provide a reliable basis for assessing whether these DIPs behave as effective TIPs. Since the reported R_0_ values seem unlikely to reflect *in vivo* reproductive capacity, caution is warranted before using these R_0_ values as evidence for or against the therapeutic viability of TIPs.

**Fig. 1.**
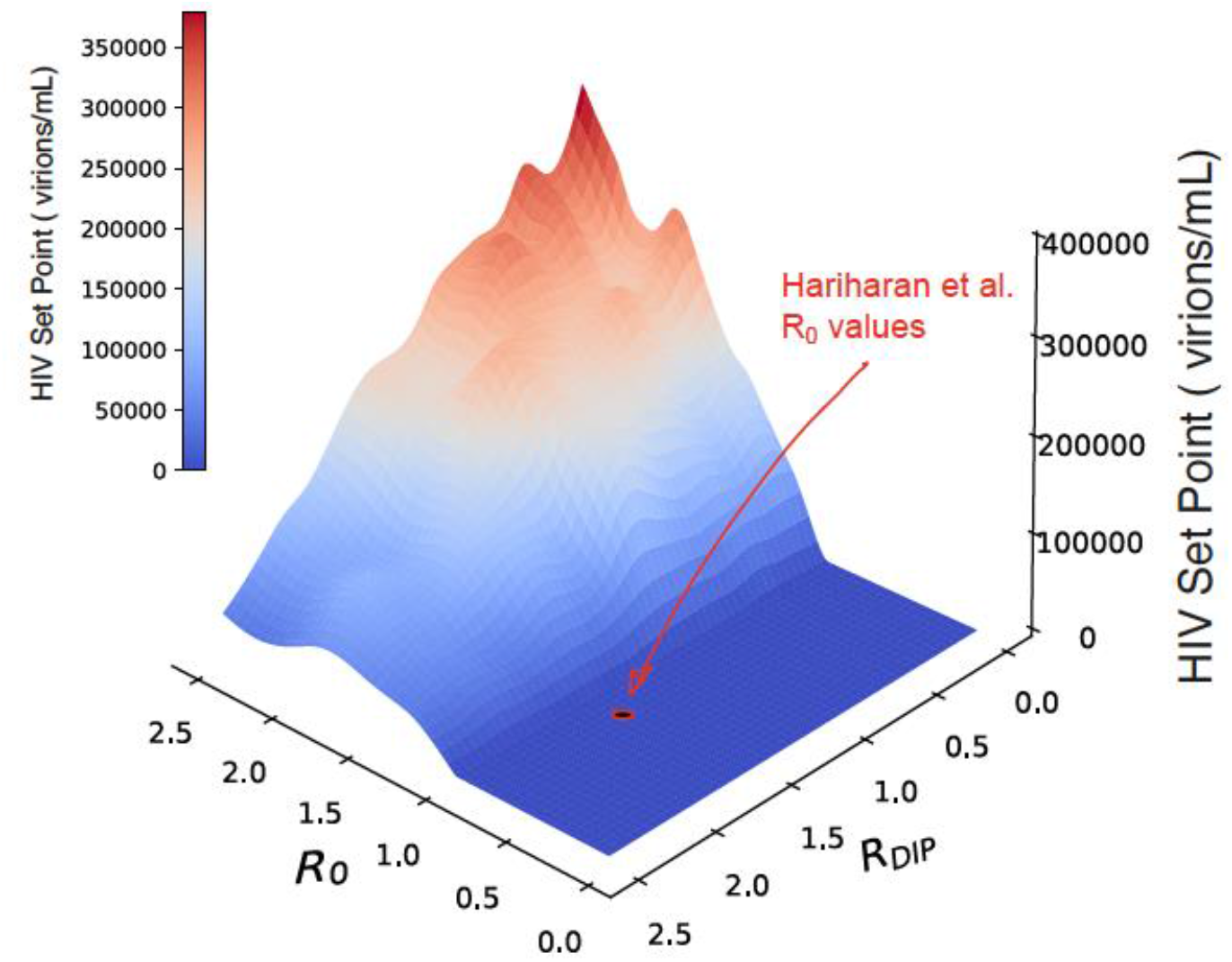
HIV set point viral load as a function of R_0_ and R_0_^DIP^. Viremia calculated from standard model of HIV-1 *in vivo* dynamics modified to include DIPs, as in Ref ^1^.

It is also difficult to reconcile how a relatively small deletion generating a Tat-deficient HIV variant could exhibit R_0_ > 1 for the reported defective genome while the corresponding full-length (non-defective) replication-competent HIV exhibits R_0_ < 1 in precisely the same assay. The reported R_0_ for the defective genome may therefore be overestimated.

Notably, the denominator used to compute defective R_0_ is inferred from reporter-positive cells at the time of co-culture—for Tat-deficient defectives, this approach excludes infected but transcriptionally silent cells that retain future reproductive potential, thereby shrinking the denominator and potentially inflating the R_0_ above unity whilst the HIV R_0_ remains below unity. Such assay-dependent artifacts further limit interpretation of these DIPs as TIP analogues.

It is also unclear why previous screens^6^ that examined constructs similar to the defective genomes isolated from P1 and P2 did not observe measurable interference, in contrast to the results reported by Hariharan et al. The interference effect of these deletions may require further study before declaring they act as TIPs and making extrapolations regarding *in vivo* efficacy of TIPs.

We note that well-established alternative mechanisms could explain the results of Hariharan et al. without invoking superinfection-mediated trafficking of defective genomes as TIP analogues.

Hariharan et al. hypothesize that in ART-treated individuals, superinfection by intact HIV enabled defective proviruses to replicate conditionally, diversify, and disseminate during suboptimal ART, generating a large heterogeneous reservoir of defective proviruses that contributes to persistent viremia and that, despite conditional replication and an estimated R_0_ > 1 in one case, these putative TIPs did not meaningfully reduce HIV pathogenesis. The data presented (Extended Data Fig. 2c-d), however, show 0% detection of the deletion during the period the authors describe as suboptimal ART (TDF/FTC/RPV), with defective genomes only becoming detectable months to years after initiation of a different optimal ART regimen. The study therefore provides no evidence that the defective genomes were present or mobilizing during suboptimal ART.

Notably, the authors do not exclude established reservoir dynamics mechanisms—e.g., reservoir ‘leakage’, the mechanism widely invoked to explain persistent wild-type HIV RNA under ART— as the source of DIP viremia without requiring ongoing superinfection. Specifically, the analysis does not consider the likelihood that clonally expanded Tat-deficient proviruses can undergo latency reversal and produce viral-like particles (VLPs), albeit at a lower level, following NF-κB– mediated activation of the LTR in the absence of Tat^7^, a phenomenon directly supported by the authors’ own data (Fig. 5g) demonstrating mobilization of Tat-deficient genomes. Lack of accessory genes does not preclude efficient VLP packaging or transduction, as 3^rd^ generation lentiviral vectors^8^ omit these genes. Thus, there is no need to invoke superinfection or complement these genes *in trans* for VLP mobilization and transduction of defective genomes. Such particles could transduce new target cells during the reported^2^ periods of suboptimal ART, obligately establish latency^9^, and expand clonally, generating the capacity for a large reservoir without requiring sustained superinfection or R_0_ > 1 spread.

Moreover, ΔVif-associated APOBEC-mediated hypermutation^10^ in the observed Vif-deleted or Vif-compromised genomes provides a well-established mechanism for the observed heterogeneity of the reservoir and sequence diversification. During the reported periods of prolonged suboptimal ART, reverse transcription during VLP-transduction events in the absence of functional Vif would be expected to generate APOBEC-edited diversity that can mimic signatures of ongoing replication without necessitating sustained superinfection-mediated spread.

## Discussion

Collectively, these well-established mechanisms may provide a more parsimonious explanation for the observations without invoking the hypothesis that the defective genomes act as TIP analogues. Thus, under this model, the observations are compatible with reservoir dynamics and do not extrapolate to TIP efficacy.

Importantly, the engineered HIV-TIP that was previously reported^1^ contains far more substantial deletions and inactivating mutations as compared to the Tat-proximal deletions in participants P1 and P2 reported by Hariharan et al.—in contrast to the Tat-proximal deletion, the HIV-TIP contains multiple deletions as well as mutations abrogating open-reading frame expression throughout the HIV genome. Consequently, the engineered HIV-TIP^1^ cannot express any HIV proteins and has no capacity to reconstitute VLPs even under maximal NF-κB or Tat stimulation, absent superinfection by HIV to provide all the missing viral proteins *in trans*. This is a fundamental difference between the TIP^1^ and the defectives reported by Hariharan et al. These biological distinctions are central when considering whether the naturally occurring defective genomes described by Hariharan et al. meaningfully model engineered therapeutic TIPs.

Importantly, none of these issues detract from Hariharan et al.’s most valuable contribution. On the contrary, by demonstrating that naturally occurring defective HIV genomes can persist without clinical harm—while underscoring that therapeutic efficacy likely requires more extensive genome excision—the study by Hariharan et al. provides strong evidence that interfering genomes are biologically relevant and tolerable in humans.

## Methods Summary

Numerical simulations utilized an established within-host virus dynamics model that is an extension of the Basic Model^11^ of virus dynamics incorporating DIP and DIP-infected populations with two additional parameters^1^. As described, the model is composed of six coupled nonlinear ordinary differential equations (Eqs. 10–15 in ref. ^1^) representing target cells, HIV productively infected cells, HIV virions, DIP-transduced cells, dually infected cells, and DIP virions. Briefly, to account for host-to-host variability and parameter uncertainty, 2000 random parameter sets were generated using a stratified Monte Carlo sampling approach (Latin Hypercube Sampling) across the eight parameters in the model, which are: *λ* (the T-cell production rate), *k* (the virus-cell infection rate), *n* (the burst size); *d* (the per-capita T-cell decay rate), *δ* (the HIV-infected cell per-capita death rate), *c* (the per-capita viral clearance rate), *P* (the intracellular increase in DIP mRNA relative to HIV mRNA in a dually infected cell), and *D* (the DIP interference parameter); *λ, P*, and *D* were sampled from uniform distributions, *k* and *n* from log-uniform distributions, and *d, δ, c* from truncated normal distributions within predefined bounds, as described^1^. For each parameter set, R_0_ and R_DIP_ were calculated as previously described^12^, and values satisfying: (i) R_DIP_ < 2.5 and R_0_ < 2.5 (in order to expedite numerical processing), and (ii) biological constraints (e.g., *δ* ≥ *d* and finite, non-negative equilibria) were retained to construct the R_0_-R_DIP_ phase space in Fig. 1. Steady states were computed analytically when *R*_*DIP*_ ≤ 1, *R*_*o*_ ≤ 1 whereas when R_DIP_ >1, steady states were integrated numerically (using stiff BDF solver, SciPy) using initial conditions *T*(0) = *λ*/*d, V*(0) = 10^2^, *V*_*t*_(0) = 10^2^, and all other states initialized to zero. The qualitative result—i.e., that no sustained viral load can be obtained when R_0_ < 1—is general^11^ and independent of the specific viral dynamics model employed.

## References

1. Pitchai, F.N.N., et al. Engineered deletions of HIV replicate conditionally to reduce disease in nonhuman primates. Science 385, eadn5866 (2024).

2. Hariharan, V., et al. Superinfection promotes replication and diversification of defective HIV-1 proviruses in people with non-suppressible viraemia. Nat Microbiol 10, 2736–2748 (2025).

3. Dahabreh, I.J. & Bibbins-Domingo, K. Causal Inference About the Effects of Interventions From Observational Studies in Medical Journals. JAMA 331, 1845–1853 (2024).

4. Boyko, E.J. Observational research--opportunities and limitations. J Diabetes Complications 27, 642–648 (2013).

5. D’Amico, F., Marmiere, M., Fonti, M., Battaglia, M. & Belletti, A. Association Does Not Mean Causation, When Observational Data Were Misinterpreted as Causal: The Observational Interpretation Fallacy. J Eval Clin Pract 31, e14288 (2025).

6. Notton, T., Glazier, J.J., Saykally, V.R., Thompson, C.E. & Weinberger, L.S. RanDeL-Seq: a High-Throughput Method to Map Viral cis- and trans-Acting Elements. mBio 12(2021).

7. Berkhout, B., Gatignol, A., Rabson, A.B. & Jeang, K.T. TAR-independent activation of the HIV-1 LTR: evidence that tat requires specific regions of the promoter. Cell 62, 757–767 (1990).

8. Dull, T., et al. A third-generation lentivirus vector with a conditional packaging system. J Virol 72, 8463–8471 (1998).

9. Razooky, B.S., Pai, A., Aull, K., Rouzine, I.M. & Weinberger, L.S. A hardwired HIV latency program. Cell 160, 990–1001 (2015).

10. Mulder, L.C., Harari, A. & Simon, V. Cytidine deamination induced HIV-1 drug resistance. Proc Natl Acad Sci U S A 105, 5501–5506 (2008).

11. Nowak, M.A. & May, R.M. Virus dynamics : mathematical principles of immunology and virology, (Oxford University Press, Oxford; New York, 2000).

12. Weinberger, L.S., Schaffer, D.V. & Arkin, A.P. Theoretical design of a gene therapy to prevent AIDS but not human immunodeficiency virus type 1 infection. J Virol 77, 10028–10036 (2003).

